# Belowground competition increases root allocation in agreement with game-theoretical predictions, but only when plants simultaneously compete aboveground

**DOI:** 10.1101/2025.01.29.635491

**Authors:** Emanuel B. Kopp, Niels P.R. Anten, Pascal A. Niklaus, Samuel E. Wuest

## Abstract

Competition among plants can lead to allocation strategies that favor individual competitiveness at the expense of group-level productivity. However, the role of root and shoot responses in driving these outcomes is intensely debated. Experimental approaches often have difficulty disentangling above- and belowground interactions due to the confounding effects of pot size and nutrient distribution. Here, we used physical dividers and varied inter-plant distances (3–24 cm) in soybean to isolate competitive interactions while controlling for these confounding effects. Simultaneous above- and belowground competition increased relative root allocation, but reduced total biomass. Aboveground competition alone had stronger effects, triggering shade avoidance and reducing both shoot and root biomass. This likely overrode belowground cues. These results underscore the significance of neighbor-induced responses, particularly under full competition, as pivotal drivers of allocation patterns and promising targets for breeding strategies that enhance collective crop performance.

## Introduction

Evolutionary game theory posits that selection in competitive environments favors resource acquisition strategies that benefit the individual but are detrimental at the level of the population (Anten & Vermeulen 2016; Gersani *et al*. 2001; Zhang *et al*. 1999). Conversely, strategies optimal for population-level productivity typically are not evolutionary stable, because such stands can be invaded by individuals that optimize their own resource acquisition at the expense of the group (Aschehoug *et al*. 2016; Harper 1977). This phenomenon is known as “tragedy of the commons” and has profound implications in agriculture, where the selection of overly competitive individuals during breeding needs to be avoided (Denison 2012; Donald 1968; Jennings & Herrera 1968; Weiner 2019).

In crops, some important traits associated with competition are well-described. Examples are plant height and an open shoot architecture. In maize, rice, and wheat, breeding for short and compact shoots accordingly contributed to substantial yield increases (Duvick *et al*. 2003; Hedden 2003; Jennings & Herrera 1968). Another trait detrimental to yield is shade avoidance, where plants sense light quality changes caused by the presence of neighbors and respond by elongating shoots and increasing leaf area, especially at the top of the canopy, where they can shade neighbors (Weiner *et al*. 2010). In soybean (*Glycine max*), shade often increases petiole and internode lengths but reduces branching (Lyu *et al*. 2023), and often leads to higher disease susceptibility and lower yield per plant (Green-Tracewicz *et al*. 2011; Lyu *et al*. 2023). Shade avoidance responses typically are undesired because they divert resources away from seed production (Pantazopoulou *et al*. 2021; Weiner *et al*. 2010). At the same time, some degree of light sensing and shade avoidance-like growth responses are required for efficient light harvesting and for canopy organization (López Pereira *et al*. 2017; Zhou *et al*. 2024). This trade-off makes targeting shade avoidance in breeding difficult.

Plants also compete belowground, but the consequences for population-level growth and yield are less well understood. Game-theoretical modelling suggests that the adaptation of plants to root competition can lead to the diversion of resources from yield to root growth, i.e. to a tragedy of the commons similar to the one observed under aboveground competition (Gersani *et al*. 2001; Zhang *et al*. 1999). Indeed, experiments by Gersani *et al*. (2001) using belowground root system separation showed that soybean plants competing belowground, i.e. without separation, had higher root biomass and lower seed yield than plants whose root systems were separated by a physical divider. However, one difficulty in interpreting this result is that pot-bounding effects may have occurred because root system separation also halved the total soil volume available to each plant (Hess & De Kroon 2007; McNickle 2020; Poorter & Sack 2012; Semchenko *et al*. 2007). An ideal experiment would simultaneously account for the effects of pot volume, total amount of nutrients per plant, and for soil nutrient concentrations, which is difficult to achieve (Chen *et al*. 2015; Maina *et al*. 2002; McNickle 2020). Recently, root-impermeable mesh dividers that allow nutrients to pass through have been used to keep available root space constant (Chen *et al*. 2021; Zhu *et al*. 2019). However, the assumption remains that root growth responses are caused by signals that pass through the mesh divider and do not require direct root contact (Cabal 2022; Chen *et al*. 2021; McNickle 2020; Zhu *et al*. 2019). Collectively, these studies show that the presence of neighbor plants can increase root biomass under belowground competition, although this effect is not always observed (Lepik *et al*. 2021; Schmid *et al*. 2015). Indeed, a recent meta-analysis confirmed such conflicting root responses in pea (Mobley *et al*. 2022).

Recently, Cabal *et al*. (2020) suggested that seemingly contradictory effects of neighbors on root growth occur because they are distance dependent. Assuming that the cost of growing and maintaining roots increases with distance from the stem, a game-theoretical model predicted that competing plants would shift additional root production to the stem because the cost of root competition is lower there. Thus, the phenomenon that emerges from such a model, which was termed “exploitative segregation of plant roots”, is one where the same resource foraging strategy leads to different root allocation responses depending on the distance between competing plants (Cabal 2022; Cabal *et al*. 2020).

Here, we studied biomass allocation in soybean plants competing aboveground, belowground, both aboveground and belowground, or not at all. We assessed root and shoot biomass at flowering, as plants had reached a size at which they strongly competed but still were growing rapidly. This allowed us to determine the competitive strategy of the plant, with organ size differences both reflecting differences in resource allocation up to this point and determining its future growth (Sun *et al*. 2021). Although this would have allowed quantification of seed yield, we chose not to assess biomass at senescence because resources would have been remobilized from the vegetative biomass, obscuring the earlier effects of biomass allocation (Liu *et al*. 2019; Maillard *et al*. 2015).

Combining the traditionally used separation treatments with a range of distances between plants (Fig. 1) allowed us to modulate the strength of competition between neighbors while keeping the available soil volume constant. Accordingly, the primary metric of interest is the statistical interaction of the separation treatment with the distance to the neighbor. Specifically, we expected neighbor-induced allocation changes to be largely absent when plants were separated or at a large distance. Under belowground competition, we expect a preferential allocation to roots, i.e. a root-to-shoot ratio that increases as plant distance decreases (Fig. 1a). Conversely, under aboveground competition, we predicted a preferential allocation to shoots, i.e. a root-to-shoot ratio that decreases as plant get closer (Fig. 1b). When plants compete both aboveground and belowground, predictions are more difficult to make because aboveground and belowground responses are coupled. We therefore expected their effects to interact. For example, the effects of belowground competition would depend on whether plants also compete aboveground (Fig. 1c).

**Figure 1:**
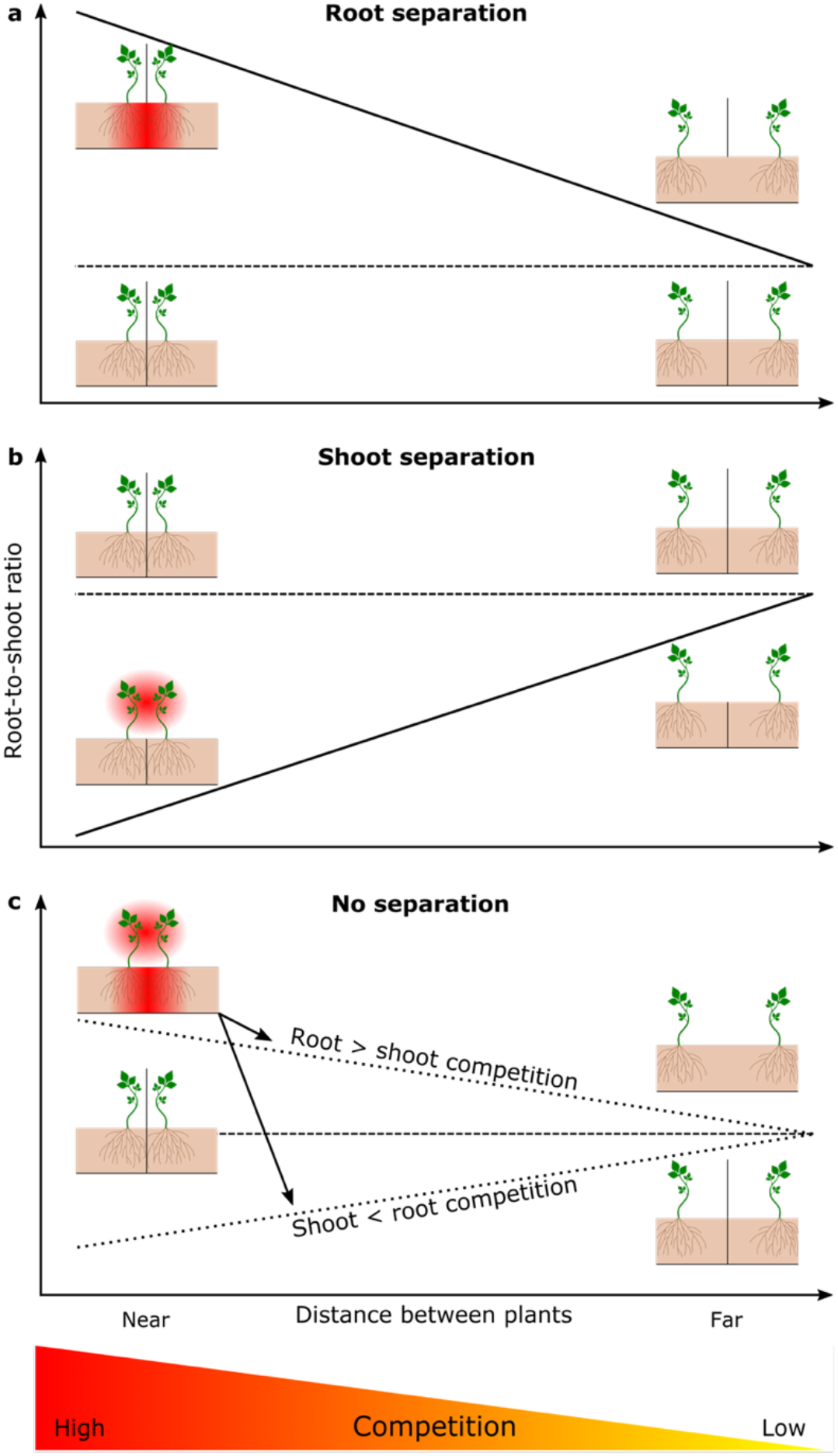
Expected ratio of root biomass to shoot biomass in dependence of root and shoot separation and inter-plant distance. Increasing inter-plant distance allows to decrease competition; hence, the effect of competition can be assessed as biomass at close distance relative to the situation when plants are far from each other. (a) When plants are always separated aboveground, we expect root competition to increase the root-to-shoot ratio. (b) When plants do not compete belowground, we expect aboveground competition to decrease the root-to-shoot ratio. (c) When plants simultaneously compete aboveground and belowground, we expect the change in root-to-shoot ratio to depend on whether the effect of competition is stronger aboveground or belowground.

## Material and Methods

In Summer 2023, an outdoor pot experiment was established in Wädenswil, Switzerland (47.222°N, 8.669°E, 509 m a.s.l.) to study competition-effects on allocation in soybean. After inoculating soybean seeds with commercial rhizobia (HiStick, BASF, Ludwigshafen, Germany; containing *Bradyrhizobium japonicum*), plants were germinated and grown in propagator trays in a greenhouse for 14 days, and seedlings then transplanted in pairs to 7.5 l rectangular pots filled with lawn soil (Ökohum Art. 2633200, Herrenhof, Switzerland) and moved outside. There, the plant pairs were separated aboveground, belowground, both aboveground and belowground, or not at all. The aboveground separation consisted of a white polystyrene sheet (80 cm high × 20 cm wide) that prevented shoot competition. The belowground separation consisted of a water-impermeable plastic sheet dividing the pot into halves. Each divider combination was combined with an inter-plant distance treatment, starting at a very close interplant distance relative to plant size, that is at a distance where plants were strongly competing (3, 6, 9, 12, 15, 18, 21, and 24 cm). Each pot contained two plants of one of four genotypes (Weber - PI 548524, Natsu Kurakake – PI 417187, Kanegawawase - PI 229330, or Kanagawa Wase - PI 506832; USDA Soybean Germplasm Collection, Urbana, IL, USA). We previously established that these four genotypes formed a tentative gradient in competitive strategies, with Weber being least and Natsu Kurakake being most competitive. Each treatment combination was replicated three times, resulting in a total of 384 pots (2 aboveground separation treatments × 2 belowground separation treatments × 8 distances × 4 genotypes × 3 replicates). Pots were protected from hail by a net and irrigated ad libitum (Figure S1 in Supporting Information).

The plants were grown for 55 days until the majority of them had started to flower. Starting on day 20, we measured plant height once a week, for a total of six measurements. On day 54, we quantified relative chlorophyll content (MultispeQ V 2.0; PhotosynQ.com; Kuhlgert *et al*. 2016). On day 55, we clipped shoots at ground level, washed roots, and dried (60°C, 48 h) and weighed all biomass samples. Because the two plants in each pot were in symmetrical positions, all biomass data were analyzed at the pot level.

We analyzed all data using linear models (aov function of R 4.3.1; http://www.r-project.org). Biomass data was analyzed without allometric correction, because the biomass data was independent of root-to-shoot ratio (R^2^ = 0.001, p > 0.5). The linear models contained row and column terms (the pots were arranged on a rectangular grid; Fig. S1) to account for spatial effects within the hail protection tunnels, followed by the full-factorial combination of aboveground and belowground dividers, the distance between the competing plants, and genotype. Given our hypothesis, we were primarily interested in the dependence of the separator effect on distance.

## Results

Genotypes differed in all traits measured (P < 0.001 for main effect of genotype, Table 1, Fig. 2). This was expected, since genotypes were deliberately chosen to represent different growth strategies in soybean. However, all other effects we were interested in were independent of genotype, i.e. there was no statistical interaction of genotype with interplant distance and aboveground or belowground separation.

**Table 1:** Statistical tests (F ratio and P-value) for effects of genotype (G), the distance between the pair of plants (D), the aboveground separation treatment (C_A_), the belowground separation treatment (C_B_), and including all interactions of the competition treatments with distance.

| Response | Explanatory variable<sup>1</sup> | F (df<sub>n</sub>, df<sub>d</sub>) | p-value |
| --- | --- | --- | --- |
| Total biomass | G | 161.03 (3, 347) | < **0.001** * |
|  | D | >0.001 (1, 347) | 0.99 |
| | D x $C_A$ | 6.845 (1, 347) | **0.009** * |
| | D x $C_B$ | 0.02 (1, 347) | 0.9 |
| | G x D x $C_A$ | 1.784 (3, 347) | 0.15 |
| | G x D x $C_B$ | 0.122 (3, 347) | 0.95 |
| | D x $C_A$ x $C_B$ | 2.62 (1, 347) | 0.106 |
| Shoot biomass | G | 132.46 (3, 347) | < **0.001** * |
|  | D | 0.14 (1, 347) | 0.71 |
| | D x $C_A$ | 5.71 (1, 347) | **0.017** * |
| | D x $C_B$ | 0.15 (1, 347) | 0.7 |
| | G x D x $C_A$ | 1.04 (3, 347) | 0.37 |
| | G x D x $C_B$ | 0.2 (3, 347) | 0.9 |
| | D x $C_A$ x $C_B$ | 5.4 (1, 347) | **0.021** * |
| Root biomass | G | 187.11 (3, 347) | < **0.001** * |
|  | D | 2.66 (1, 347) | 0.6 |
| | D x $C_A$ | 5.49 (1, 347) | **0.02** * |
| | D x $C_B$ | 0.69 (1, 347) | 0.79 |
| | G x D x $C_A$ | 2.2 (3, 347) | 0.088 |
| | G x D x $C_B$ | 0.05 (3, 347) | 0.99 |
| | D x $C_A$ x $C_B$ | 0.09 (1, 347) | 0.76 |
| Root-to-shoot ratio | G | 164.88 (3, 347) | < **0.001** * |
|  | D | 0.793 (1, 347) | 0.37 |
| | D x $C_A$ | 0.006 (1, 347) | 0.94 |
| | D x $C_B$ | 0.58 (1, 347) | 0.45 |
| | G x D x $C_A$ | 0.786 (3, 347) | 0.5 |
| | G x D x $C_B$ | 0.172 (3, 347) | 0.92 |
| | D x $C_A$ x $C_B$ | 4.163 (1, 347) | **0.042** * |
| Final height | G | 136.85 (3, 347) | < **0.001** * |
|  | D | >0.001 (1, 347) | 0.98 |
| | D x $C_A$ | 2.24 (1, 347) | 0.135 |
| | D x $C_B$ | 0.611 (1, 347) | 0.435 |
| | G x D x $C_A$ | 0.113 (3, 347) | 0.952 |
| | G x D x $C_B$ | 0.3 (3, 347) | 0.825 |
| | D x $C_A$ x $C_B$ | 4.774 (1, 347) | **0.03** * |
| Shoot-biomass-to-height ratio | G | 67.55 (3, 347) | < **0.001** * |
|  | D | 0.11 (1, 347) | 0.74 |
| | D x $C_A$ | 4.031 (1, 347) | **0.045** * |
|  | D x C<sub>B</sub> | 0.01 (1, 347) | 0.908 |
|  | G x D x C<sub>A</sub> | 1.101 (3, 347) | 0.349 |
|  | G x D x C<sub>B</sub> | 0.992 (3, 347) | 0.397 |
|  | D x C<sub>A</sub> x C<sub>B</sub> | 2.176 (1, 347) | 0.141 |
| Relative chlorophyll content | G | 6.545 (3, 347) | < **0.001** * |
|  | D | 2.98 (1, 347) | 0.085 |
|  | D x C<sub>A</sub> | 1.761 (1, 347) | 0.185 |
|  | D x C<sub>B</sub> | 0.244 (1, 347) | 0.621 |
|  | G x D x C<sub>A</sub> | 1.282 (3, 347) | 0.28 |
|  | G x D x C<sub>B</sub> | 0.696 (3, 347) | 0.555 |
|  | D x C<sub>A</sub> x C<sub>B</sub> | 4.06 (1, 347) | **0.045** * |

<sup>1</sup>: To avoid confounding with pot volume, competition treatments can only be interpreted in interaction with inter-plant distance.

**Figure 2:**
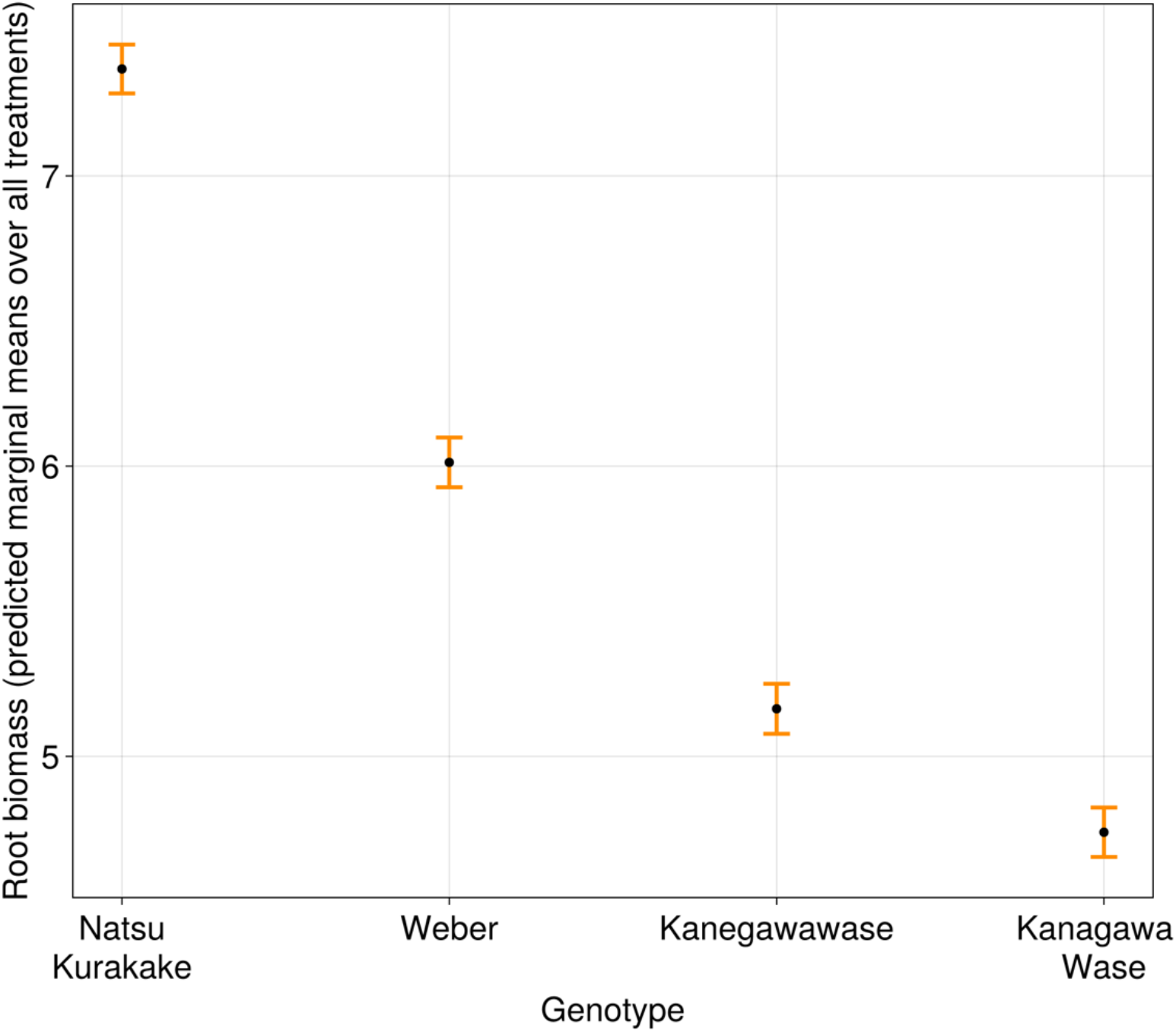
The four genotypes strongly differed in all measured traits, as shown here for root biomass. Error bars indicate +/– one standard error of the mean.

Our primary interest was on effects of root competition. Some aspects of plant growth, however, depended only on shoot separation. First, shoot competition caused shade avoidance responses. This manifested in lower shoot-biomass-to-height ratios, i.e. plants were more slender when close together (aboveground separation x distance F_1,347_ = 4.03, P < 0.05; Table 1, Fig. 3a). Second, both root and total biomass decreased under shoot competition (aboveground separation x distance: F_1,347_ > 5 and P < 0.05 for both; Table 1, Fig. 3b, c).

**Figure 3:**
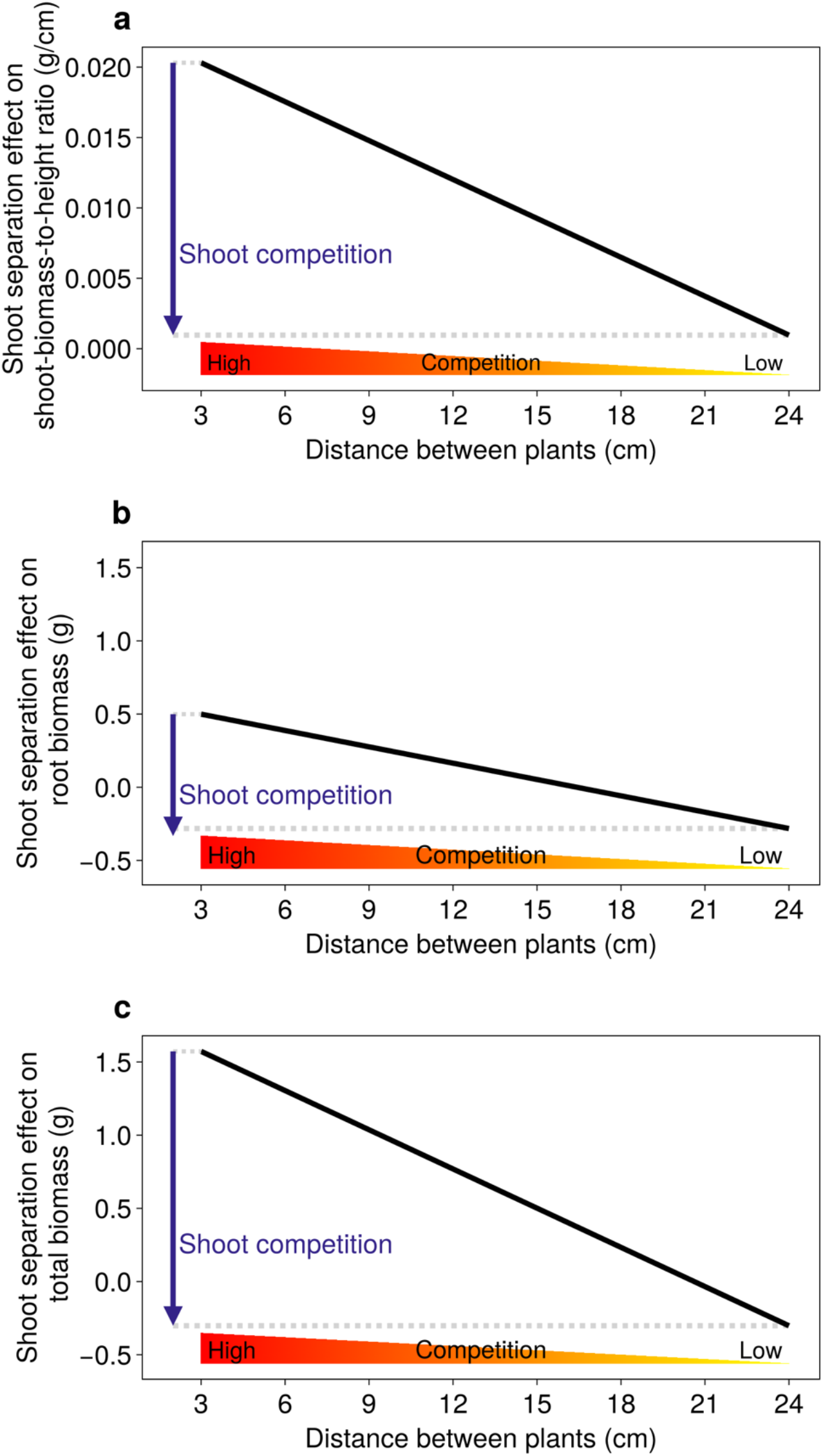
Shoot competition effects on (a) shoot biomass to height ratio, (b) root biomass, and (c) total biomass. The shoot separation effect is the difference between pots with and without shoot separation. We used the distance treatment to modulate the strength of competition. Hence, shoot competition manifests as difference between the separation effects when plants are close and when plants are distant. Lines show model predicted values.

All other growth-related parameters also depended on root separation. Specifically, root competition effects were contingent on shoot competition, manifesting as three-way interactions between the two separation treatments and distance. In the absence of aboveground separators, the effects of root competition were consistent with our hypothesis. Specifically, root-to-shoot ratios increased under belowground competition, and this effect was strongest when plants were close together. In contrast, no such increase was found when plants were separated aboveground. These opposing patterns manifested as statistical three-way interaction (aboveground separation x belowground separation x distance, F_1,347_ = 4.16, P < 0.05; Fig. 4a). Conversely, shoot competition decreased root-to-shoot ratios, as expected, but only when plants were separated belowground (Fig. 4b). Similar contingencies of root competition effects on shoot competition and vice-versa were found for final plant height and relative chlorophyll content (aboveground separation x belowground separation x distance, F_1,347_ > 4, P-values < 0.05; Table 1; Fig. S2). For example, plants competing above- and belowground were smaller than plants competing belowground only, and plants had higher relative chlorophyll content when competing belowground but not aboveground.

**Figure 4:**
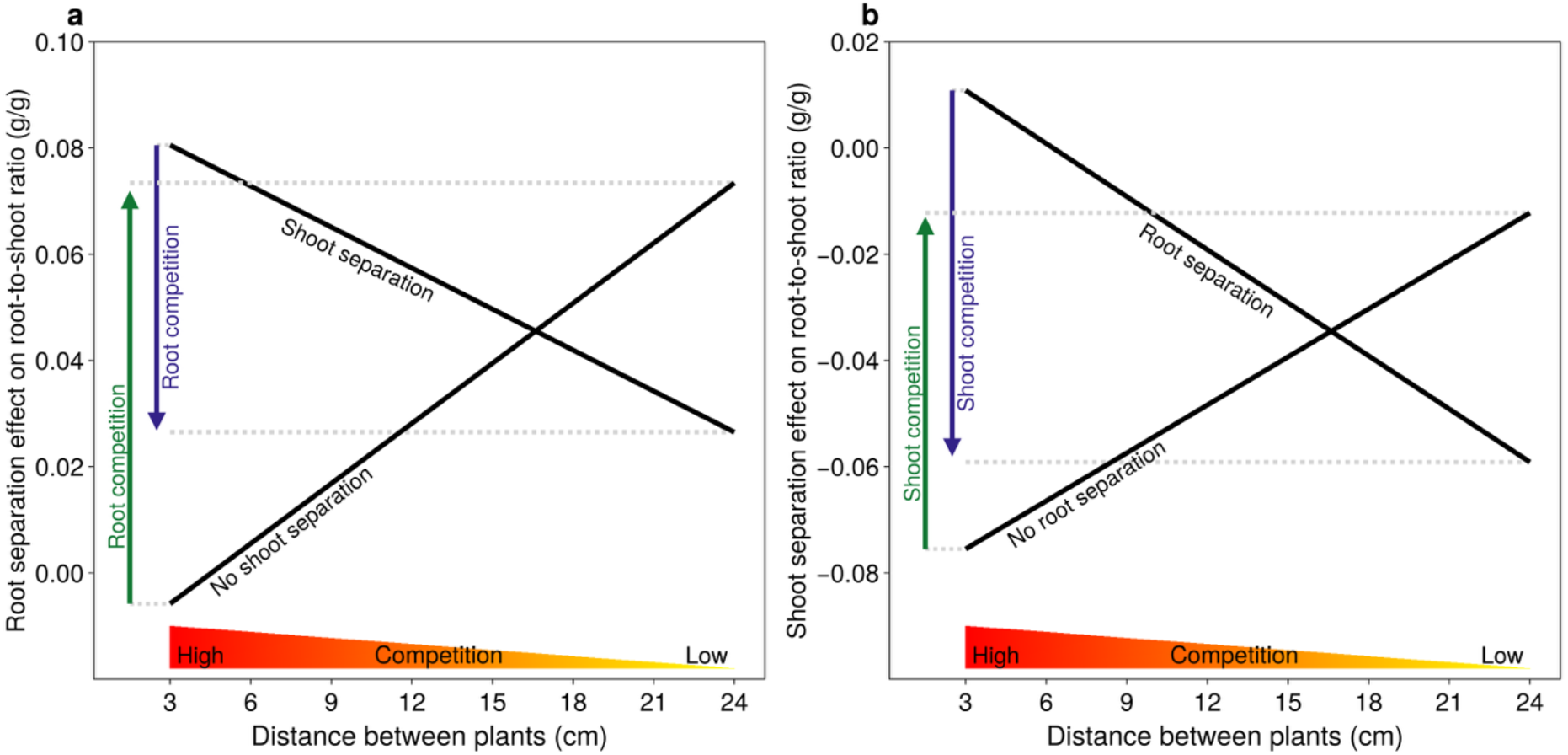
(a) Root separation effects on root-to-shoot ratio, in dependence of inter-plant distance. When plants were not separated aboveground, root competition increased root-to-shoot ratio (green arrow). However, when plants were separated aboveground, root competition decreased root-to-shoot ratio (blue arrow). (b) Shoot separation effects on root-to-shoot ratio. When plants were not separated belowground, shoot competition increased root-to-shoot ratio (green arrow). However, when plants were separated belowground, shoot competition decreased root-to-shoot ratio (blue arrow). The dependency of root competition effects on shoot separation manifested as statistical interaction between the two separation treatments and distance (F_1,347_ = 4.16, P < 0.05, Table 1).

Shifts in root-to-shoot ratio may be caused by changes in root biomass, in shoot biomass, or in both, and we therefore examined what caused the significant three-way interaction of treatments and interplant distance on root-to-shoot ratios in our experiment. Belowground competition had contrasting effects on shoot biomass, depending on whether plant competed aboveground (aboveground separation x belowground separation x distance, F_1,347_ = 5.4, P < 0.05, Fig. 5). This increased root-to-shoot ratio when plants were not separated aboveground, and reduced root-to-shoot ratios when they were. In contrast, root competition did not affect root biomass. However, shoot competition decreased root biomass (Fig. 3b), a change which lowers root-to-shoot ratio. Overall, our results indicate that the observed three-way interaction in root-to-shoot ratio is predominantly driven by changes in shoot biomass.

**Figure 5:**
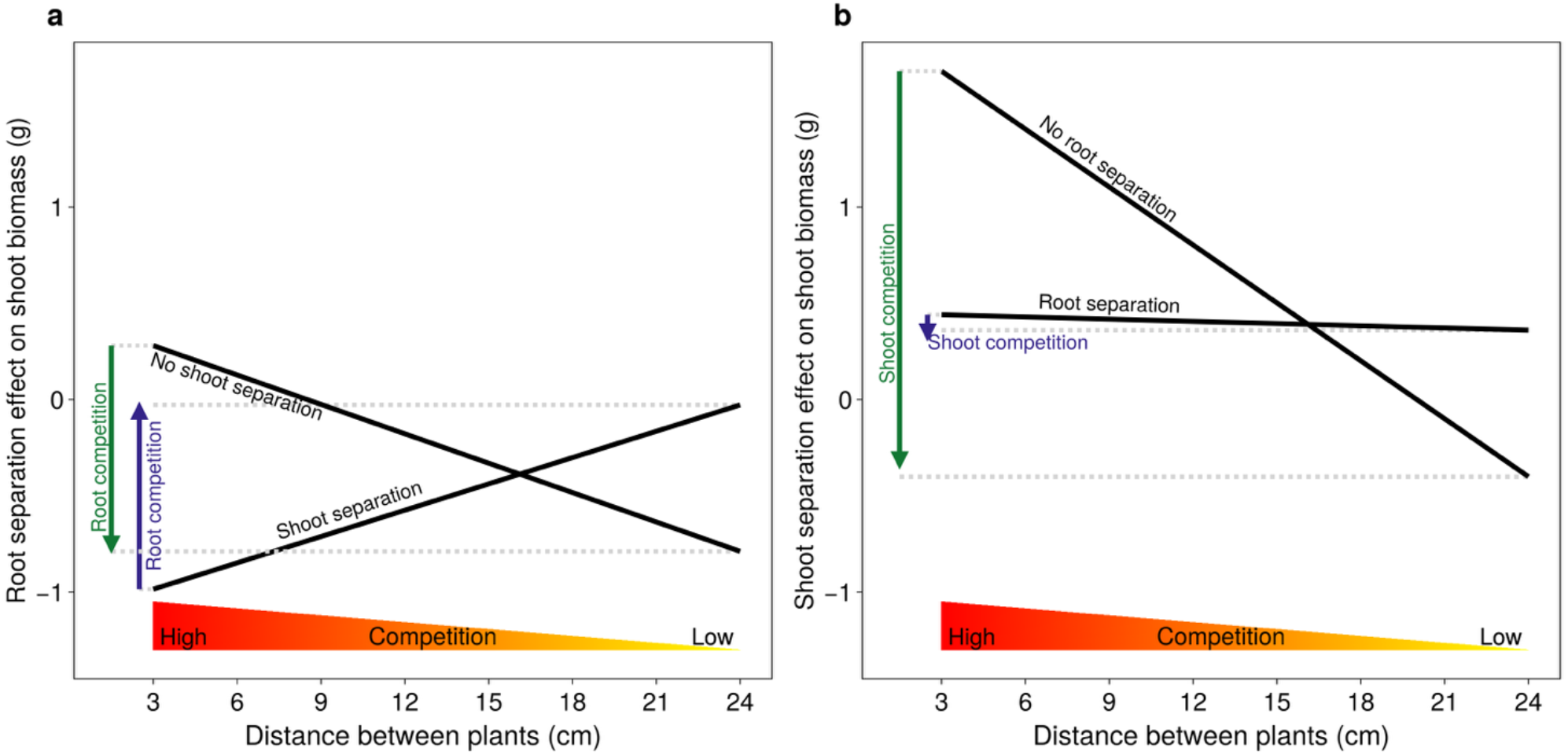
(a) Root separation effects on shoot biomass, in dependence of inter-plant distance. When plants were not separated aboveground, root competition decreased shoot biomass (green arrow). When plants were separated aboveground, root competition increased shoot biomass (blue arrow). (b) Shoot separation effects on shoot biomass, in dependence of inter-plant distance. When plants were not separated belowground, shoot competition decreased shoot biomass (green arrow). When plants were separated belowground, shoot competition slightly decreased shoot biomass (blue arrow). The dependency of root competition effects on shoot separation manifested as statistical interaction between the two separation treatments and distance (F_1,347_ = 5.4, P < 0.05, Table 1).

## Discussion

### Concomitant aboveground and belowground competition shifts allocation to roots

In our experiment, we found increased relative allocation to roots and decreased total biomass when plants were close to their neighbors, but only when plants were simultaneously competing aboveground and belowground. In contrast, aboveground competition led to a reduction of total biomass, irrespective of whether plants also competed belowground. Aboveground competition also reduced final root biomass, which also was independent of belowground competition. Our data thus indicates increased allocation to root growth under full competition, but nevertheless a lower final root biomass. We think that occurs because increased allocation to root competition reduces allocation to photosynthetically-active leaf area and thus slows the plants’ overall growth rates, which translates into a lower final root biomass because plants become smaller. This pattern is in line with a tragedy of the commons in which competition with neighbors reduces the combined performance of the two plants. Our results are in line with Gersani *et al*. (2001), who manipulated belowground competition using physical dividers and found evidence for such a tragedy of the commons for seed production (Hardin 1968). It is interesting to note that Gersani *et al*. (2001) used a setup where competing plants were grown relatively close to each other, without aboveground dividers and in a greenhouse over the winter period requiring supporting artificial light – conditions that typically induce a shade avoidance response. This growth condition was similar to our close-proximity and no-aboveground-separator treatment, and the observed increased root allocation under such conditions is consistent with our study. We therefore think that many discrepancies among studies could be due to differences in experimental conditions such as light levels and light quality, interplant distances, or pot sizes (Murphy & Dudley 2007). Particularly interplant distance, which in our experiment interacted strongly with both above- and belowground competition, has received surprisingly little attention in past work, despite its potential to modulate competition intensity and neighbor detection (Cabal *et al*. 2020; Gottlieb & Gruntman 2022; McNickle 2020; Semchenko *et al*. 2007).

### Shade avoidance responses under aboveground competition led to a stronger decrease in shoot biomass than in root biomass

Responses to aboveground competition were consistent with the typical shade avoidance syndrome (Ballaré & Pierik 2017; Green-Tracewicz *et al*. 2011; Pantazopoulou *et al*. 2021), i.e. plants were more slender and produced less root biomass. Plant height typically increases under shade avoidance (Pierik & De Wit 2014). However, under shoot competition alone, final plant height increased only slightly. On the contrary, simultaneous shoot and root competition had a clear negative effect on final height. This reduction in height likely resulted from a reduction in total plant biomass due to less-optimal allocation under shade avoidance. We think that this is induced through light quality changes when neighbors are present, and that there was no limitation of growth by available light (Fig. S1). Even though shade avoidance can increase the fitness of individuals at high densities, our results and those of others suggest that the light-foraging strategy invoked by the presence of neighbors is detrimental to the performance of the overall stand (Pantazopoulou *et al*. 2021; Schmitt *et al*. 1995; Wille *et al*. 2017).

### Aboveground competition has stronger effects than belowground competition

Competition between two plants ranges from fully symmetric to strongly asymmetric: depending on the extent to which larger plants acquire amounts of resources that are proportionately or disproportionately large relative to their size (Weiner 1990). Light competition is usually asymmetric, because the taller plant can pre-empt light, as light is a largely directional resource. Nutrient competition is more symmetric with nutrient acquisition largely proportional to the size of the root system (Schwinning & Weiner 1998). Based on our findings, we think that aboveground competition was stronger than belowground competition. Indeed, the interaction between aboveground separation and neighbor distance was significant for multiple measurements. However, the interaction between belowground separation and neighbor distance was never significant in absence of an interaction with aboveground separation. An integration of above- and belowground responses to competition on root growth could occur through various processes. For example, plants may “sense” the strength of a neighboring competitor aboveground and adjust their belowground foraging strategy (Gottlieb & Gruntman 2022). Alternatively, as predicted by game theoretical models for belowground interactions, aboveground interactions with neighbors may increase the costs of root production and maintenance (Cabal *et al*. 2020).

### The response to neighbors is a valuable breeding target

Here, we showed a novel and simple way to control for the confounding factors that have long dominated the debate on how to interpret competition experiments involving pairs of plants. We strongly suggest that future work investigating the effect of individual foraging strategies on plant group performance should explicitly account for neighbor distance. It has long been anticipated, most famously by Donald (1968), that reducing competitive traits in crop plants may hold great potential for increasing yields (see also Duvick *et al*. 2003; Jennings & Herrera 1968). However, this has only recently become a renewed focus in the emerging research field of “crop evolutionary ecology” (Anten & Vermeulen 2016; Denison 2012; Weiner 2019; Weiner *et al*. 2017). Here, we found reduced biomass production under shoot competition, despite similar resource availability. We also found a diversion of resources to root production under combined root and shoot competition. We quantified plant biomass at flowering to capture effects of allocation when plants were still rapidly growing, and found a tragedy of the commons with respect to total plant biomass. It is plausible that a similar tragedy of the commons would have been found for seed yield if the experiment had been run for longer (but see Murphy & Dudley 2007). The fact that we found this effect across a range of genotypes that varied substantially in allocation patterns suggest that this finding is general. Overall, this suggests that responses to the presence of neighbors bear potential as breeding target.

## Acknowledgements

This work was supported by the Swiss National Science Foundation, project 310030_192537. We acknowledge Lukas Vonmetz and Jürgen Krauss (both Agroscope) for technical support.

## Competing interests

The authors declare no competing interests.

### Author contributions

E.B.K., S.E.W. and P.A.N conceptualized and designed the research (with input from N.P.R.A.). E.B.K and S.E.W performed the experiments. E.B.K performed the analyses and wrote the manuscript with input from S.E.W and P.A.N. All authors revised and approved the final version of the manuscript.

### Data accessibility statement

The data that support the findings of this study are openly available on Zenodo at http://doi.org/10.5281/zenodo.15524543

## Supporting Information

**Figure S1:**
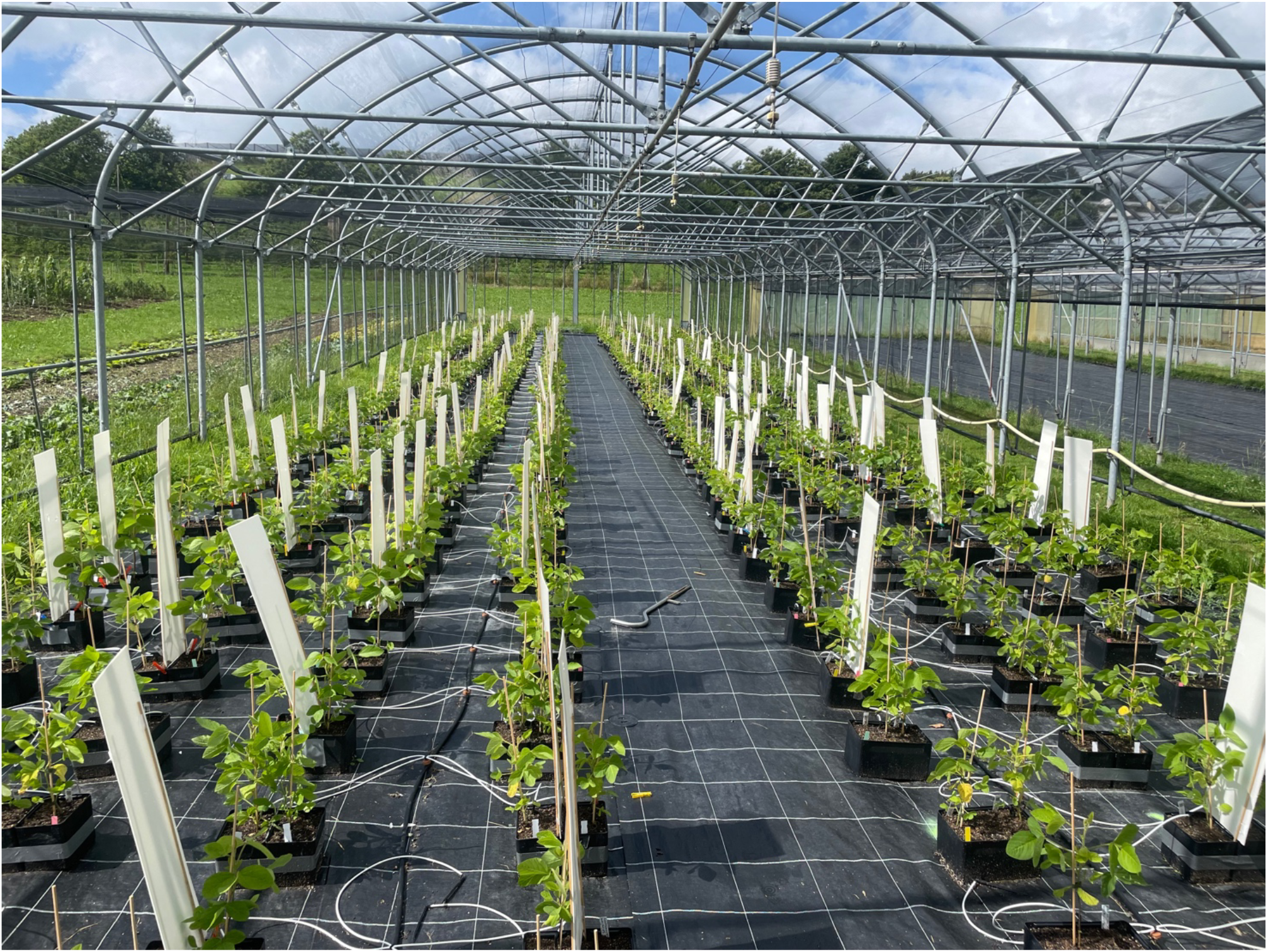
Experiment under hail protection net, at 41 days after seed germination. The conditions were chosen so that water or light limitations were excluded as much as possible.

**Figure S2:**
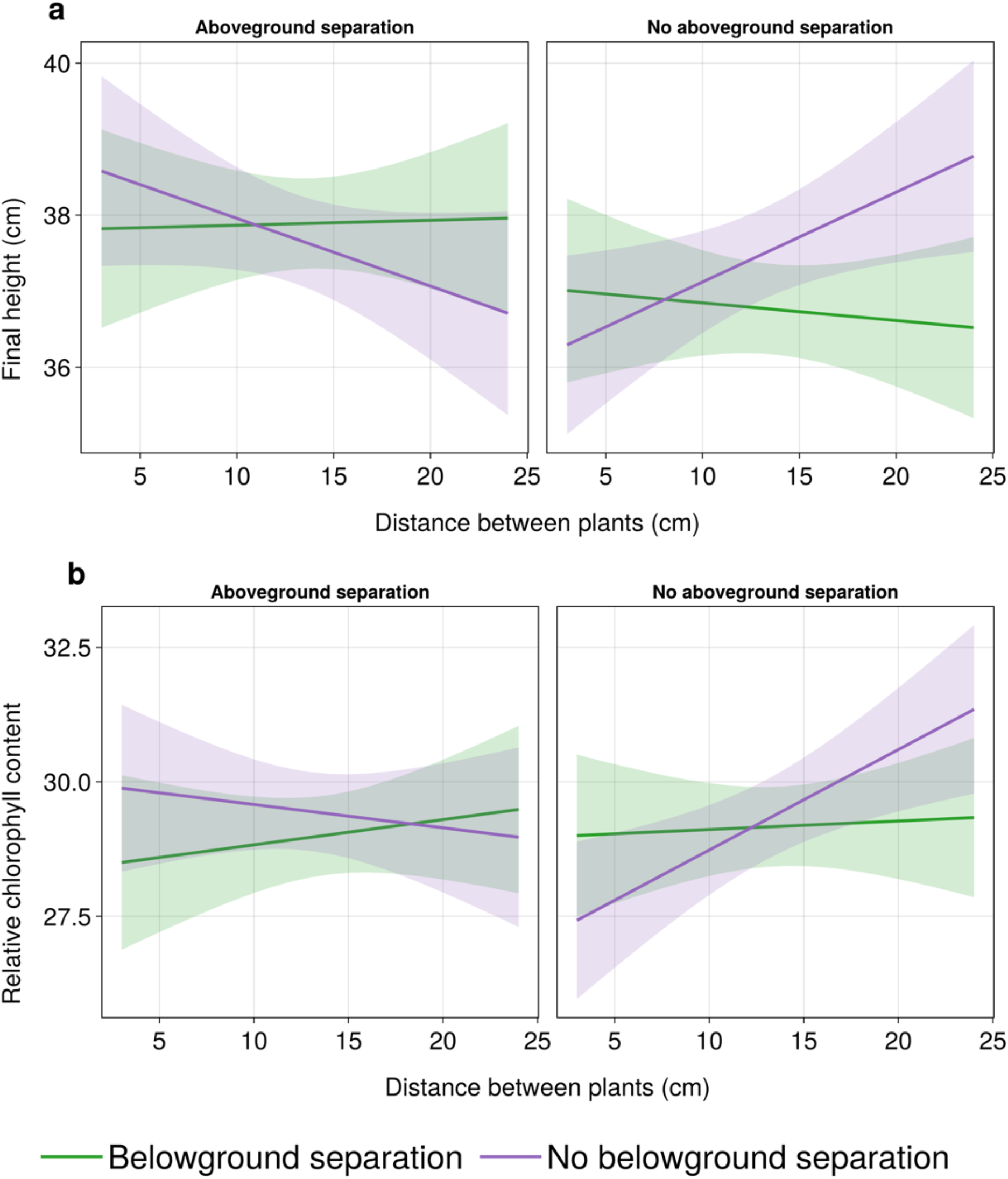
Predicted marginal means for (a) final plant height, and (b) relative chlorophyll content depending on presence/absence of aboveground separation (panels) and on presence/absence of belowground separation (colors). The opposing patterns of the lines between the panels show the three-way interactions between the two separation treatments and the interplant distance.

